# Immune defense in *Drosophila melanogaster* depends on diet, sex and mating status

**DOI:** 10.1101/2022.05.01.490225

**Authors:** Kshama Ekanath Rai, Han Yin, Arnie Lynn C. Bengo, Madison Cheek, Robert Courville, Elnaz Bagheri, Reza Ramezan, Sam Behseta, Parvin Shahrestani

## Abstract

Post-mating immunosuppression has been widely accepted as a female trait in *Drosophila melanogaster*. Our results challenge this notion by presenting a mating-immunity trade-off in males as well as in females. When inoculated with the fungal pathogen, *Beauveria bassiana*, both males and females die faster compared to inoculated virgins, and survival is lower when inoculated flies are continuously mated compared to a single day of mating. Past studies with *Beauveria bassiana* have shown females to be more susceptible to infection than males. Our results challenge this finding as well, showing that the direction of sexual dimorphism in immune defense depends on mating status, specific *Beauveria bassiana* strain, and fly genotype. Moreover, we show that survival after fungal infection is largely influenced by diet, and that post-infection dietary improvements can help enhance survival. Post-mating suppression in *Drosophila* survival of *B. bassiana* infection presents study opportunities with potential applications for biological control of insect vectors of human disease and insect crop pests.

## Introduction

Mosquito disease vectors can be catalysts for the spread of malaria, dengue fever, and other devastating diseases within human populations (1,2). The entomopathogen *Beauveria bassiana*, which grows naturally in soil and infects a broad range of insect hosts, has been demonstrated to have potential uses against mosquito disease vectors as well as insect crop pests that threaten food security (1,3,4), and bed bugs (5).

*Drosophila melanogaster*, the laboratory fruit fly, has an immune system similar to that of insect disease vectors and pests, contributing to its value as a model organism.

Moreover, the *D. melanogaster* immune system has many similarities to the mammalian innate immune system (6). Similar to human skin, the fly cuticle provides a physical barrier of defense against pathogens. Cellular and humoral responses of the innate immune system further help ward off infection (7, 8, 9). The ease of maintenance and short life cycle of *D. melanogaster* make it an optimal candidate for experiments involving many individuals and multiple replicates.

*D. melanogaster* exposure to *B. bassiana* in the wild generally happens on the cuticle surface. Upon contact, spores germinate and penetrate the cuticle, growing as hyphae inside the host, and later sporulating on the cadaver (10). Female *D. melanogaster* are more susceptible than males to five tested strains of *B. bassiana* (11). The results of Shahrestani et al. (11) suggest that sexual dimorphism in defense is, at least in part, controlled by humoral defenses. Specifically, the sexual dimorphism is ablated by mutations in the Toll pathway, which is known to be involved in fighting Gram positive bacteria and fungi, and also by mutation of the gene *Relish*, which is in the IMD pathway thought to be involved in fighting Gram negative bacteria (11). Further improving our understanding of sexual dimorphism in insect defense against *B. bassiana* could aid in the development of effective biological control efforts that can target reproductive female insects.

Sexual dimorphism in post-infection survival of *D. melanogaster* has also been seen with the bacterial pathogens *Providencia rettger*i and *Providencia alcalifaciens* (12). With these pathogens, not only are females more susceptible than males, but mated females are more susceptible than virgin females (12), with inoculated virgin females surviving better than inoculated mated females and carrying lower bacterial loads (12). Given this mating-immunity trade-off, a potential explanation for the increased susceptibility to infection in females relative to males is that females allocate more resources to reproduction (13). One explanation is that females enter a post-mating state, which is induced by receiving sex peptide from male seminal fluid (14). In these studies, the matingimmunity trade-off is thought to be a female-specific trait. As such, virgin females are thought to have more male-like immune defense (12). But, it is possible that both males and females experience a mating-immunity trade-off. Indeed, McKean et al. (15) showed evidence for a trade-off between male sexual activity and male immunity.

Here we show that both females and males exhibit a tradeoff between mating and post-infection survival and that continuous mating further reduces survival. Post-infection survival was sexually dimorphic, but the direction of this dimorphism depended on mating status. Moreover, we show that the direction of sexual dimorphism in post-infection survival additionally depends on fly genotype, fungal strain, and diet. Thus, we uncover more of the complexity of post-mating immune suppression and sexual dimorphism in immune defense.

## Materials and Methods

### Drosophila melanogaster Population

We used genetically diverse *D. melanogaster* populations, specifically the C3 population of Shahrestani et al. (16). Population C3 is maintained with non-overlapping 14-day generations at population size of 2,000 flies at ∼25° C, 12h light:12h dark cycles. For the past 72 generations, Population C3 was maintained at the California State University in Fullerton on a Cornmeal diet (10.0 g/L agar, 24.2 g/L yeast, 59.2 g/L cornmeal and 76.7 g/L molasses), and prior to that it was maintained at Cornell University on a Glucose diet (100g/L yeast, 100g/L glucose, 1% Drosophila agar). At the start of each experiment, flies were reared in vials at densities of 60-80 eggs per vial as in our other work (16).

### Fungal Pathogen

As in Shahrestani et al. 2021, two strains of the entomopathogenic fungus *Beauveria bassiana* were used: GHA, which was obtained from Mycotech, Inc. (now Bioworks, Inc., Victor NY, lot number TGA1-96-06B), and ARSEF 12460 (Shahrestani and Vandenberg; USDA Agricultural Research Service Collection of Entomopathogenic Fungi, Ithaca, NY). Fungal spores were stored at -4° C. Prior to use, spores were allowed to warm to room temperature.

### Fungal Inoculation

Fungal inoculations were done as previously described (11, 16). Specifically, flies were briefly anesthetized with CO_2_ and then moved onto Petri dishes placed on ice to maintain anesthesia while they were sprayed either with 5mL of a control fungal-free suspension of 0.03% Silwet (Plant media, a division of bio world) in autoclaved DI water, or with 5mL of a fungal suspension of 0.3g of *Beauveria bassiana* spores (either ARSEF 12460 or GHA) suspended in 25 mL of 0.03% Silwet. Sprays were done using a custombuilt spray tower (17). The spray dose was double checked by placing a microscope slide cover next to the flies during inoculation, then resuspending the slide and counting the number of spores in 10uL suspensions. Each spray introduced ∼10^3^ spores/mm^2^ onto the surface of anesthetized flies. After the spray, flies from each treatment were moved to separate acrylic cages (volume: 450 cm^3^), fed with a Petri dish of fly medium, and kept at ∼100% humidity for 24 hours. In high humidity conditions, fungal conidia germinate, and the hyphae penetrate the insect cuticle, entering the hemocoel (18). After 24 hours, humidity was reduced to 60% for the duration of each assay.

### Mortality

Dead flies were removed daily from all cages to avoid secondary inoculation of live flies by spores on the cuticles of the deceased flies and to record the numbers of dead females and males. Food plates were replaced with fresh ones daily.

### Experiment 1: Effects of mating and sex on post-infection survival of *D*. *melanogaster* when inoculated with *B*. *bassiana* strain ARESEF 12460

To test the effects of mating on post-infection survival, three mating conditions (virgin, mated, and cohabit) were established as follows. To ensure virginity of flies, on day five from egg, third instar larvae were transferred from rearing vials into individual straws using a paint brush. The straws had Cornmeal food on one end and were sealed with pipette tips on both ends. On day 12 from egg (2-3 days post eclosion), emerged flies were sexed while still in the straws, and the food in the straw was replaced. On day 16 from egg, flies were moved from straws to vials at densities of 30 flies/vial, using brief CO_2_ anesthesia. At this stage, the 30 flies in the vial were either all male or all female for the virgin groups, or half male and half female for the mated and cohabiting groups. After 24 hours, on day 17 from egg, the flies were sprayed with fungus or control suspension. For each spray, 60 flies were anesthetized using carbon dioxide for 5 minutes on a CO_2_ pad to allow sorting by sex, then placed on ice for the ∼2 minutes of the spray time. For the virgin groups, males and females were sprayed separately and kept in separate cages after the spray. For the mated groups, after 24 hours of mating in vials, males and females were sprayed separately and maintained in separate cages. Lastly, for the cohabiting groups, after 24 hours of mating in vials, males and females were sprayed together and cohabited in the same cages after the spray. Flies were in cages at densities of 60 same sex flies per cage, or 30 males and 30 females per cage. At least 257 flies per sex per treatment were tested. Food plates in cages were replaced daily, and eggs laid on the plates by cohabiting and 24-hour mated females were kept and incubated for 7 days until pupation, at which point the number of pupae were counted as a proxy of offspring count. Mortality and fecundity were followed for 21 days post spray. The entire experiment was replicated four times.

### Statistical Analysis of Experiment 1

All analyses were performed in the statistical software R (https://www.r-project.org/). To determine the effects of mating on the survival of female and male *D. melanogaster* when inoculated with the entomopathogenic fungus, *B. bassiana*, we used the Proportional Hazard model (19) with a novel approach via Bootstrap for creating confidence intervals for survival probabilities. Before building the model, we tested the difference among all the replicates by Kaplan-Meier survival function (20) as well as log-rank test (21) and found no differences among the four replicates. To check the proportional hazards assumption, a scaled Schoenfeld residual was plotted and a test using the Schoenfeld residuals against the transformed time was conducted for each covariate. Because some covariates broke the assumption, a piecewise PH model was proposed, in which the time splitting points were selected by a stratified model. With additional interaction terms, a piecewise PH model was chosen using both likelihood test score and Akaike information criterion (22). To detect any influential outliers, every observation was assessed by its delta-beta value for each predictor of the best model. Influential outliers were kept in the model as we had no reason to think they resulted from error. The PH model not only allows for hazard ratio estimates but also for predicted survival probabilities. We resampled observations of each fly group 1,000 times with replacement to construct 95% bootstrap intervals for a linear combination of the covariates in the model.

To quantify the effect of mating on survival percent in the study, a piecewise Proportional Hazard model was proposed (Model 1).

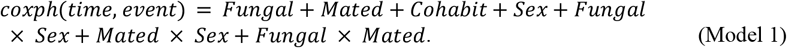

In this piecewise PH model, the 21-day experiment time was split into three time intervals: 0-5, 5-11, and 11-21. This transformed the covariates Fungal, Mated, and Cohabit to time-dependent ones. Each time-dependent covariate has different coefficient estimates over the time intervals:

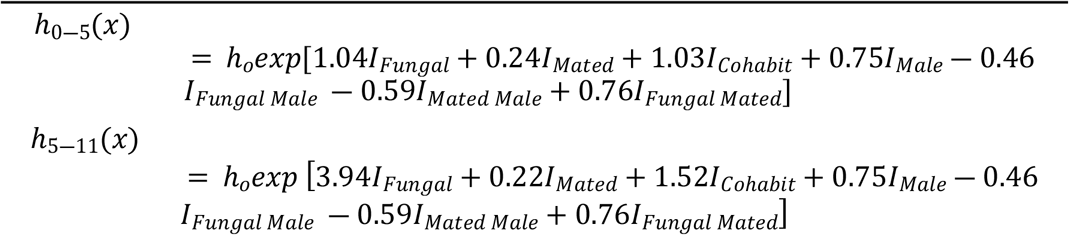

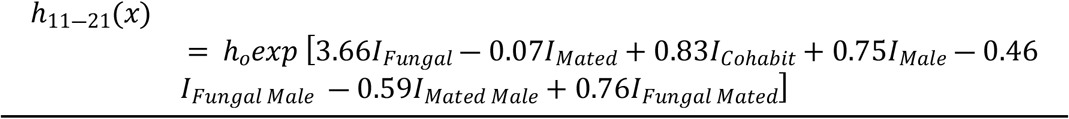

The piecewise PH model gave us a measure of the effect of mating on survival at any time point and allowed for further visual inspection of the survival probabilities over the 21 days of the experiment. We also investigated the effects of infection and mating status on average offspring count per surviving female via the analysis of variance (ANOVA) (23). We first calculated the number of offspring counts per surviving female fly and assessed the mean values of different groups of treatments and mating statuses. The analysis not only examined the main effects but also explored the interaction effects between treatments and mating statuses. Based on the final model, the means with standard errors and 95% confidence intervals were calculated. Also, post-hoc comparisons were performed to indicate which groups were significantly different from others.

### Experiment 2: Sexual dimorphism in post-infection survival of *D*. *melanogaster* when inoculated with *B*. *bassiana* strain GHA

On day 12 from egg, flies were transferred out of rearing vials and into fresh vials in groups of 15 males and 15 females. To separate and count the flies by sex, we anesthetized them using a CO_2_ gun for 10 seconds and put them on the CO_2_ pad for 3 minutes. On day 17 from egg, flies were inoculated with GHA or control suspensions. For each spray, 60 flies were anesthetized with CO_2_ for 15 seconds to immobilize them and placed on Petri dishes on ice for the ∼2 minutes of the spray time. Males and females were sprayed together and cohabited together after the spray. Mortality was followed for 21 days post spray. At least 500 flies were tested per sex per treatment. The experiment was replicated four times.

### Statistical analysis of Experiment 2

Sexual dimorphism in survival after GHA inoculation was analyzed similarly as in Experiment 1 (see Model 2).

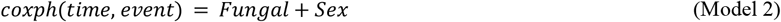

The time-dependent covariates have different coefficient estimates over the time intervals 0-8 days, 8-12 days and 12-21 days:

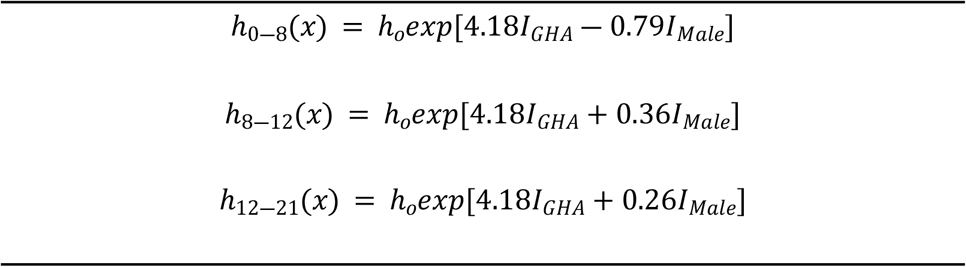

### Experiment 3: Effects of Cornmeal and Glucose diets on sexual dimorphism in postinfection survival of *D*. *melanogaster* inoculated with *B*. *bassiana* strain GHA

On Day 12 from the egg, flies were transferred, in groups of 15 males and 15 females, to fresh vials containing an assigned diet: cornmeal or glucose (ingredients provided above). To separate and count the flies by sex, we anesthetized them using a CO_2_ gun for 10 seconds and put them on a CO_2_ pad for 3 minutes. On day 15 from egg, the flies were sprayed with either GHA and control suspensions. Sprayed flies were moved to cages and fed with either the cornmeal or glucose diets. The before and after spray diet combinations are summarized in Table S1. Approximately 150 flies were sprayed per condition and kept in plexiglass cages (volume = 450 cm^3^). Dead flies were removed and sexed daily for 12 days (until age 27 days from egg). The experiment was replicated three times.

### Statistical Analysis of Experiment 3

Sexual dimorphism in survival after GHA inoculation and different diet conditions was analyzed similarly as in Experiment 1. A piecewise PH model was proposed (see Model 3).

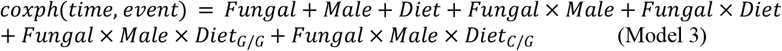

In this piecewise PH model, the 12-days after the spray (the days for which there is mortality data) were split into three time intervals: 0-3, 3-9, and 9-12. Each time-dependent covariate has coefficient estimates over these time intervals. The baseline *h*_*o*_ is females without infection. The estimated hazard ratio (*h*_*t*_(*x*)/ *ho*) at a specific time point *t* between any other group *x* and the baseline can be obtained by the coefficient estimates:

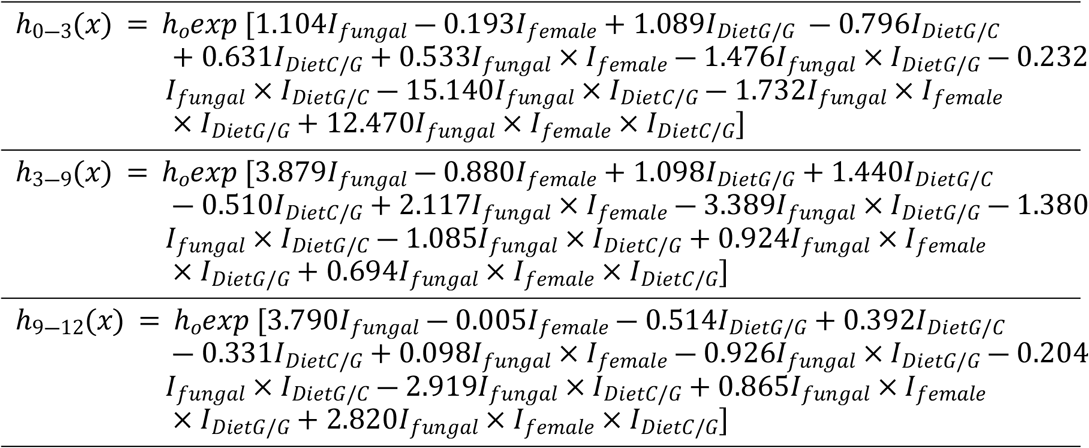

### Experiment 4: Effects of yeast supplementation on sexual dimorphism in post-infection survival of *D*. *melanogaster* inoculated with *B*. *bassiana* strain GHA

On Day 12 from egg, 15 males and 15 females were transferred to a fresh vial containing an assigned diet: cornmeal, glucose, or cornmeal with yeast supplement. The Cornmeal plus yeast supplement diet was made by adding a layer of yeast paste to the surface of the Cornmeal diet. The yeast paste was made by using a ration of 1g of yeast to 5 mL of DI H_2_O. Then using a pipette, 5mg of yeast was added onto the surface of the Cornmeal food in each vial, and 80mg of yeast was added onto the Cornmeal food in each Petri dish, which gives the same amount of yeast per surface area for vials and plates.

Flies were handled in the same way as in Experiment 3. The experiment was replicated four times. The dietary conditions of Experiment 4 are shown in Table S2.

### Statistical Analysis of Experiment 4

Sexual dimorphism in survival after GHA inoculation and different diet conditions was analyzed similarly as in Experiment 1. A piecewise PH model was proposed (see Model 4).

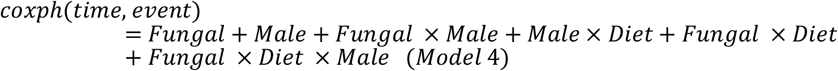

The model was split into three time intervals: 0-4 days, 4-9 days, and 9-12 days after spray. Over these time intervals, the coefficient estimates were different:

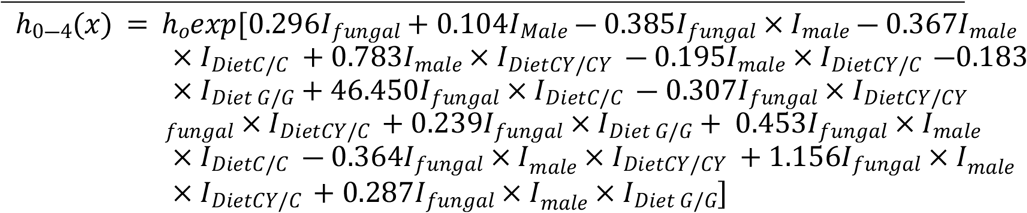

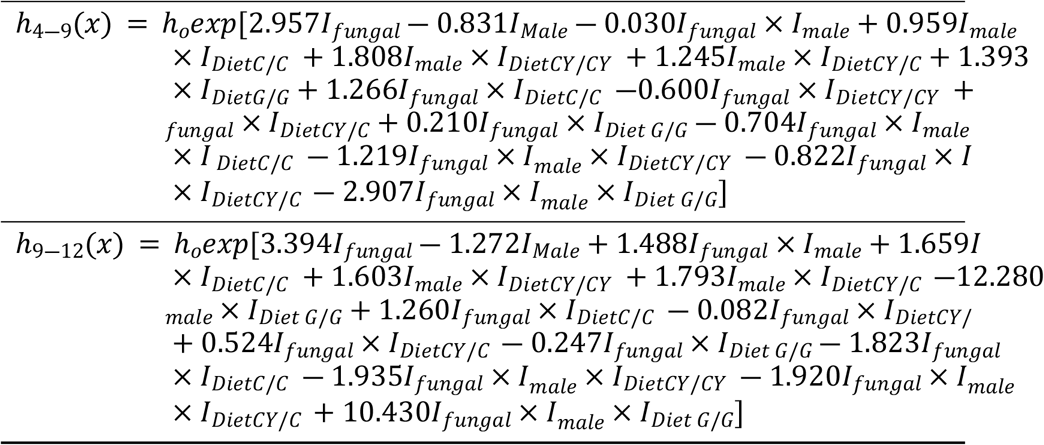

### Experiment 5: Effects of varying levels of yeast supplementation on sexual dimorphism in post-infection survival of *D*. *melanogaster* inoculated with *B*. *bassiana* strain GHA

Flies were reared on Cornmeal diet, then on day 12 from egg, they were moved to fresh vials containing Cornmeal diet at densities of 15 male and 15 females per vial. On day 15, flies were sprayed as in Experiment 3. After the spray, they were moved to cages and fed with Petri Dishes of food that differed in their level of yeast supplementation. One condition used the same supplementation as in Experiment 4 (C/CY1.0), a second condition used 0.5x the amount of yeast (C/CY0.5), and a third condition used 1.5x the amount of yeast (C/CY1.5). A control condition used Cornmeal diet without supplementation (C/C). The yeast levels were adjusted by altering the amount of yeast suspended in DI water, such that the same total volume of suspension was added to the food dishes.

### Statistical analysis of Experiment 5

Sexual dimorphism in survival after GHA inoculation and different diet conditions was analyzed similarly as in Experiment 1. A piecewise PH model was proposed (see Model 5).

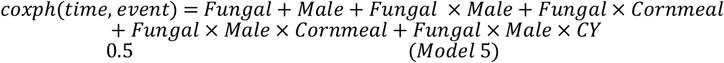

The model was split into three time intervals: 0-5 days, 5-8 days, and 8-12 days after spray. Each time interval has a model formula given below. Over these time intervals, the coefficient estimates were different:

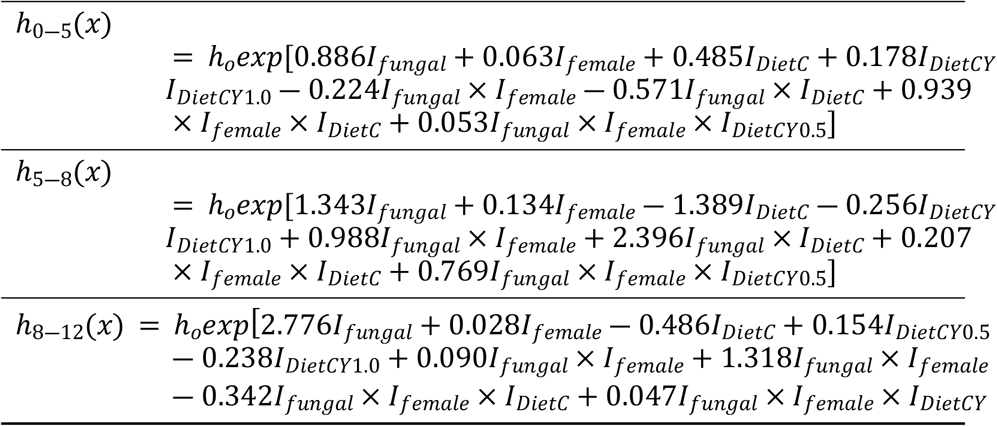

## Results

### Mating status affects survival after inoculation with *B*. *bassiana* ARSEF 12460 in both male and female *D*. *melanogaster* (Experiment 1)

Inoculated flies died faster than control flies, suggesting the effectiveness of *B. bassiana* infection (Fig 1). Among infected females, virgins survived better than mated and cohabiting females in all time intervals (Fig 1, Fig S1, Table S3), and mated females survived better than cohabiting females (Fig 1, Fig S1, Table S3). For males, the same pattern was true in the 5-11 and 11-21 day intervals (Fig 1, Fig S1, Table S3). Even among control (uninfected) females and males, virgins showed a significantly higher survival than cohabiting flies in the 5-11 and 11-21 intervals (Fig 1, Fig S1, Table S3).

**Fig 1.**
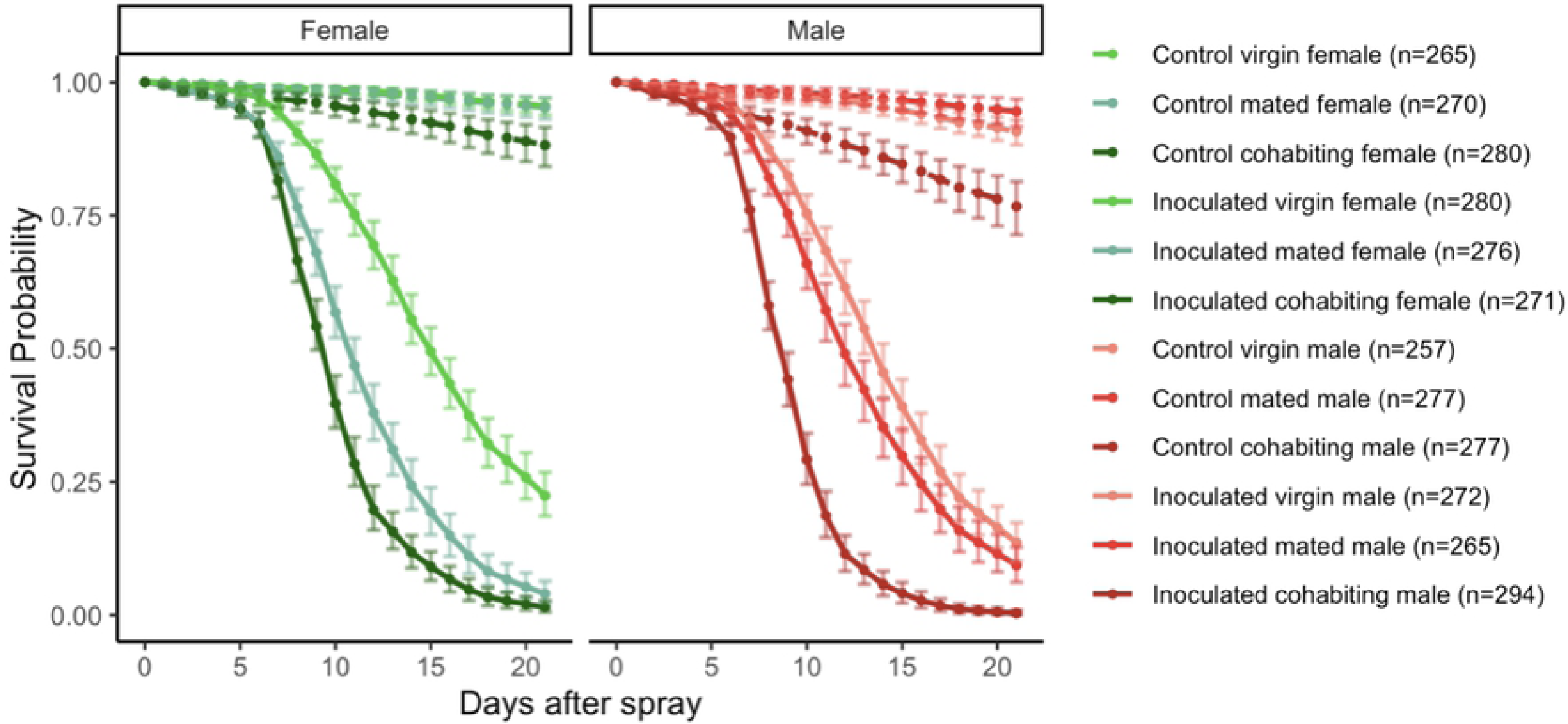
Survival post inoculation with *Beauveria bassiana* is affected by mating status in both males and females. Data from Experiment 1. Flies were sprayed with fungal suspension (*n* = 1658, solid lines) or with a control, fungus-free suspension (*n* = 1626, dashed lines) at age 17 days from egg, and survival was followed for 21 days. In both females (green) and males (red), mated flies (which mated for 24 hours prior to the spray) had lower survival than virgin flies, and flies that mated for longer than one day (cohabiting flies, which mated for 24 hours prior to spray and then cohabited with the other sex after the spray) had lower survival than mated flies (see Table S1 for statistical analysis). This figure shows model estimates for survival proportions, which are obtained from the raw survival data (Figure S1).

### Survival after inoculation with *B*. *bassiana* ARSEF 12460 is sexually dimorphic, but the direction of the dimorphism depends on mating status (Experiment 1)

Sexual dimorphism in post infection survival was observed in all three mating statuses. Among infected cohabiting flies, females showed a better survival than males (*p*-value = 0.013, Fig S1, Table S4). The same trend was observed in virgin flies (*p*value = 0.0016, Fig S1, Table S4). However, the trend was reversed in mated flies, where the males survived better than females (*p*-value = 0.0084; Fig S1, Table S4).

### Reproductive output is affected by infection and mating status (Experiment 1)

Mated and cohabiting females had offspring counts that declined with age over the span of 21 days post inoculation (Fig 2A). As the flies aged, the fungal inoculated groups showed lower offspring counts than the controls, as indicated by the *p*-value of <0.001 in the interaction between days and treatment (Table S5). In the control groups, females that cohabited with males had more offspring than females that mated only for one day, but in the fungal inoculated groups, the cohabiting and single-day mated females had the same numbers of offspring measured by counting pupal casing (Fig 2B).

**Fig 2.**
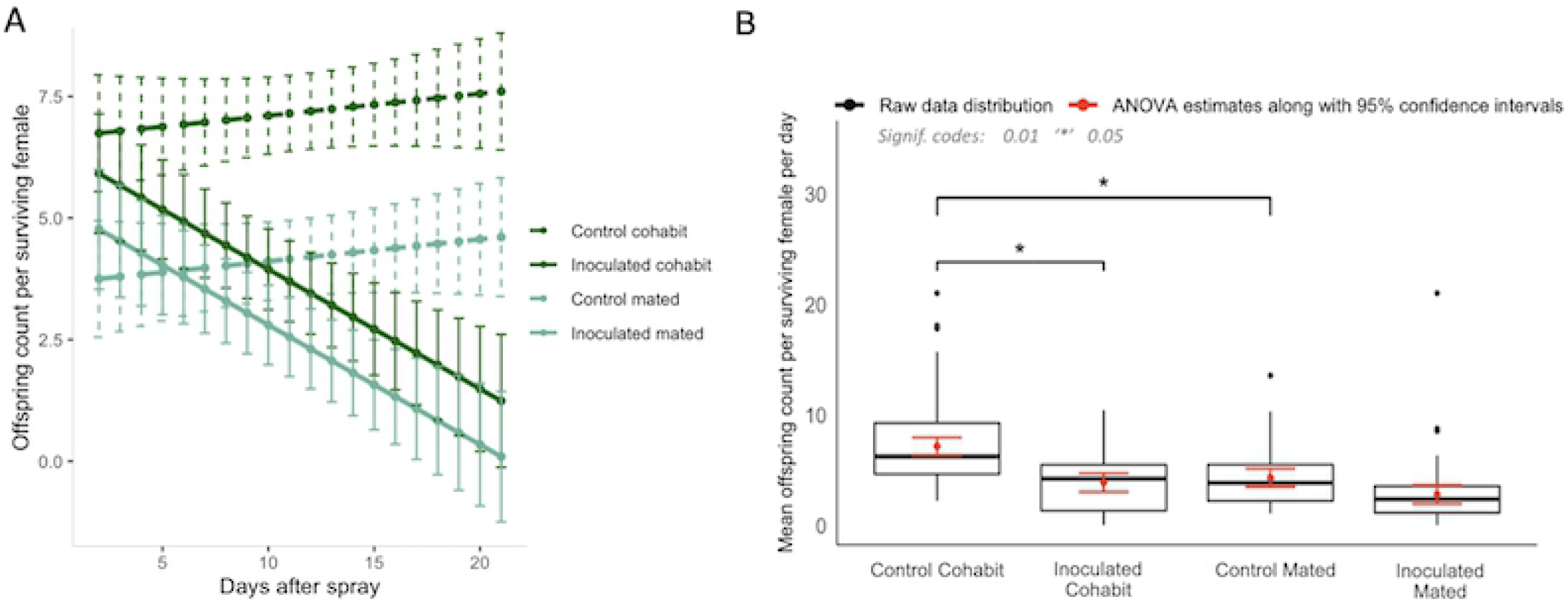
Reproductive output of mated and cohabiting females from both control and fungal inoculated conditions. (A) Analysis of variance for the offspring counts produced by control (dashed lines) and fungal inoculated (solid lines) females at two different mating statuses. The data points are plotted for 95% confidence intervals. The fungal inoculated groups showed a lower offspring count in comparison to the control groups (p-value : 3.313×10^−8^). The mated groups showed a lower offspring count in comparison to the cohabit group in control treatment (p-value: 4.534×10^−6^). There was an interaction effect between treatment (fungal/control) and mating status (cohabit/mated) with a p-value of 3.725×10^−2^. (B) Box plot shows the raw data distribution of offspring counts per surviving female and predicted mean value for cohabit and mated flies. Black dots represent the outliers while red lines represent the 95% confidence intervals. Control cohabiting females have a higher offspring count than the fungal cohabiting females (p-value: 0.01203) whereas there was no significant difference between offspring counts in control mated and fungal mated groups (p-value: 0.08702). Control cohabiting females had a significantly higher offspring count than the control mated (p-value: 0.01687). However, inoculated cohabiting and inoculated mated groups did not show a significant difference in offspring counts (p-value: 0.17707). The control cohabiting females (predicted mean : 7.124) have a lower offspring counts than fungal cohabiting females (predicted mean: 3.844) and the same was seen in control mated (predicted mean : 4.275982) and fungal mated females (predicted mean : 2.772).

### Survival after inoculation with *B*. *bassiana* GHA is sexually dimorphic, but only for the first few days after inoculation (Experiment 2)

Shahrestani et al. (11) reported that when *D. melanogaster* Population C3 (same outbred population used here) was inoculated with GHA under cohabiting conditions, males survived better than females in the ten days post-inoculation. Their study did not extend beyond 10 days. Replicating their study, we found that after inoculation with GHA, males survived better than females for the first 12 days after inoculation, but thereafter the female and male survivals converged for the remainder of our 21-day study (Fig 3, Table S6).

**Fig 3.**
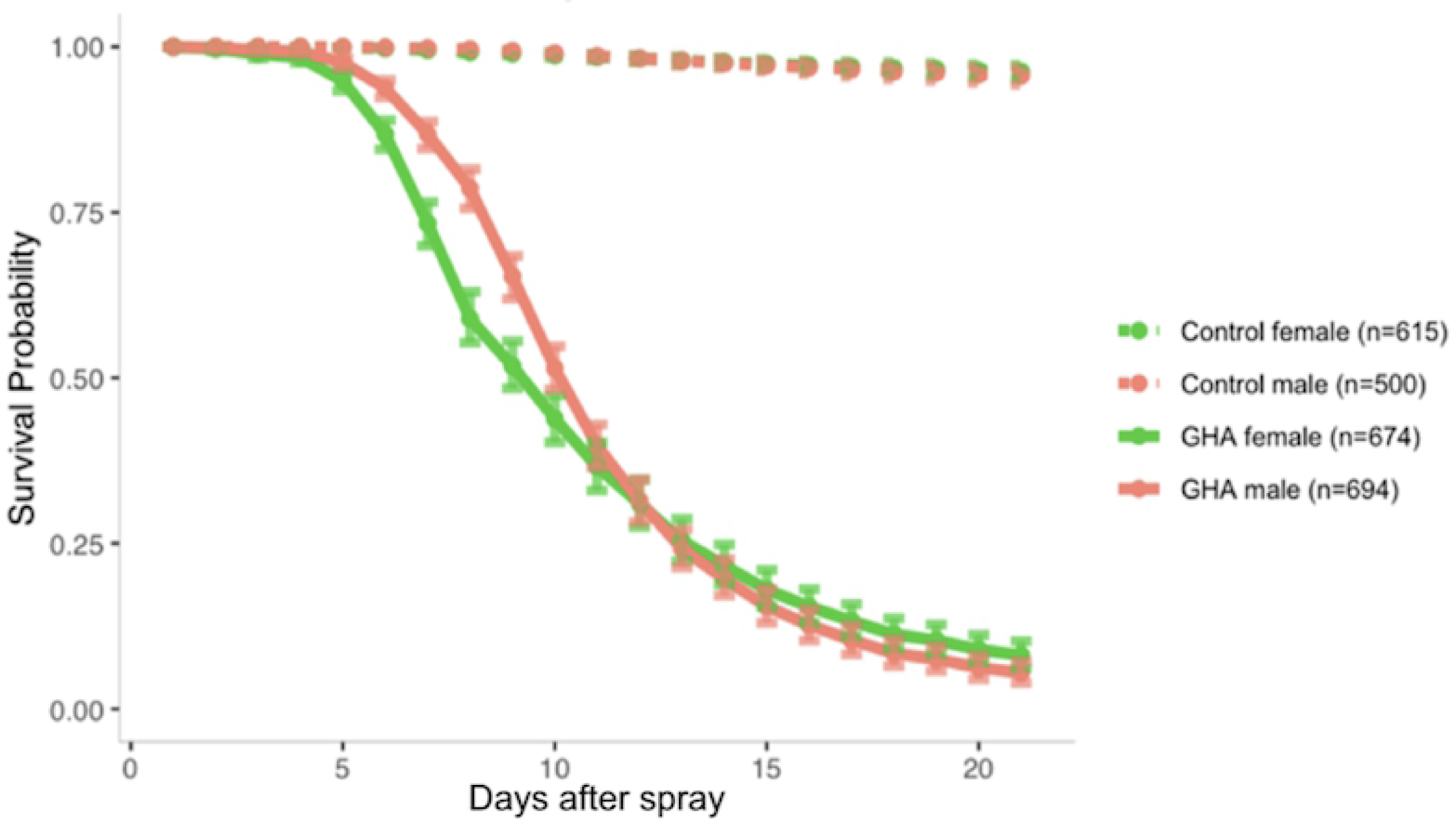
Sexual dimorphism in survival of *D. melanogaster* after inoculation with *B. bassiana* strain GHA. Data from Experiment 2. Figure shows model estimates with 95% Bootstrap confidence intervals from the raw data shown in Figure S2. Inoculated males (Orange) survived better than inoculated females (Green) until day 12 post spray. See Table S4 for statistical analysis.

### Introduction of Glucose diet pre- and postinoculation with *B*. *bassiana* GHA affects survival in sex-specific ways (Experiment 3)

Control flies survived better than inoculated flies regardless of diet (Fig 4). Flies that received Glucose diets post-spray survived better after inoculation with *B. bassiana* (Fig 4). The time of introduction of the Glucose diet impacted the survival of flies, predominantly in male flies who survived better than females (Fig 4). Table S7 summarizes the statistical comparisons of hazard ratios among males and females of control and inoculated groups under the various dietary conditions. Among uninfected controls, there was no sexual dimorphism in survival observed under any dietary condition. Among inoculated flies, when the post-inoculation diet was Cornmeal, regardless of whether the pre-inoculation diet was Cornmeal or Glucose, females had higher hazard than males (survived worse) in the 3-9 day interval post inoculation, but male and female hazards were the same in the 0-3 and 9-12 day intervals post inoculation (Table S7). When flies were given a Glucose diet post inoculation, regardless of whether the pre-inoculation diet was Cornmeal or Glucose, males survived better than females in both the 3-9 and 9-12 day intervals post inoculation (Table S7). Table S8 summarizes the effects of diet on survival in control and inoculated flies relative to survival on Cornmeal diets. In both females and males, receiving a Glucose diet improved post-inoculation survival in most of the time intervals (Table S8).

**Fig 4.**
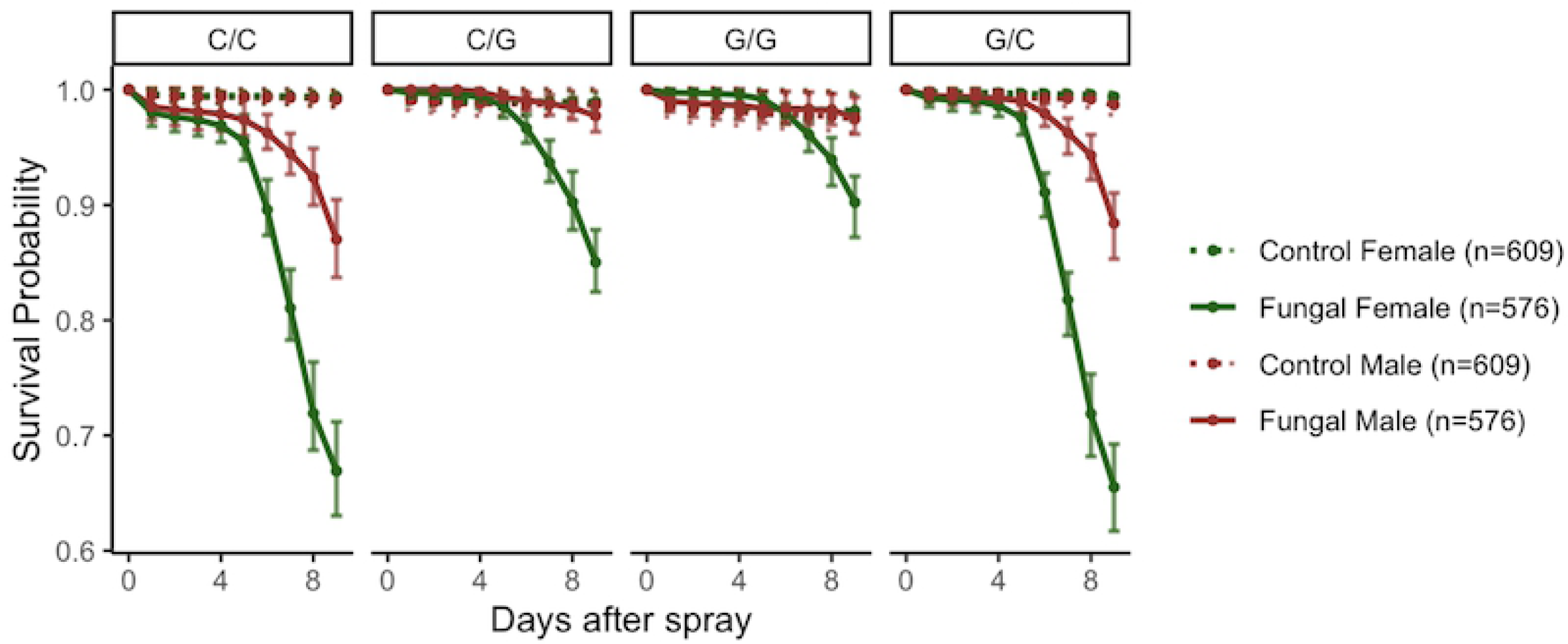
Effects of diet on post-inoculation survival of *D. melanogaster* inoculated with *B. bassiana* strain GHA. Data from Experiment 3. Figure shows model estimates with 95% Bootstrap confidence intervals from the raw data shown in Figure S3. For analysis of these data see Tables S4 and S5. Inoculated flies (Solid lines) had lower survival than control flies (Dashed lines). There was no sexual dimorphism observed among control flies, but among inoculated flies males (Red) survived better than females (Green). The timing of introduction of a glucose diet affected post inoculation survival.

### Yeast supplementation pre- and postinoculation with *B*. *bassiana* GHA positively impacts survival, but not as much as the Glucose diet (Experiment 4)

In all dietary conditions, control males and females showed a higher survival than the fungal inoculated males and females (Fig 5). Cornmeal fed flies who received Cornmeal diet before and after the spray had the lowest post-inoculation survival (Fig 5). Flies that were fed with cornmeal before spray, but yeast supplemented cornmeal after the spray showed a rescue in survival in both males and females (Fig 5, Table S9). Flies fed with yeast supplemented cornmeal before the spray and cornmeal after the spray showed lower survival than flies that received yeast supplementation before and after inoculation (Fig 5, Table S9). The highest survival was seen in flies that were given Glucose diets before and after the spray, followed by the condition in which flies received yeast supplemented cornmeal diet before and after the spray (Fig 5). Yeast supplementation affected sexual dimorphism in surviving inoculation (Fig 5, Table S10). When yeast supplementation was provided after inoculation, it ablated the sexual dimorphism in survival of inoculation (Table S10). However, when yeast supplementation was provided before inoculation, but not after, sexual dimorphism in survival was present in the 3-9 day time intervals, similarly to the control Cornmeal diet (Table S10). On the Glucose diet, males survived better than females in both the 3-9 day and 9-12 day time intervals (Table S10). Under uninfected conditions, when yeast supplementation was provided before and after the spray, male survival was negatively impacted relative to female survival (Table S10).

**Fig 5.**
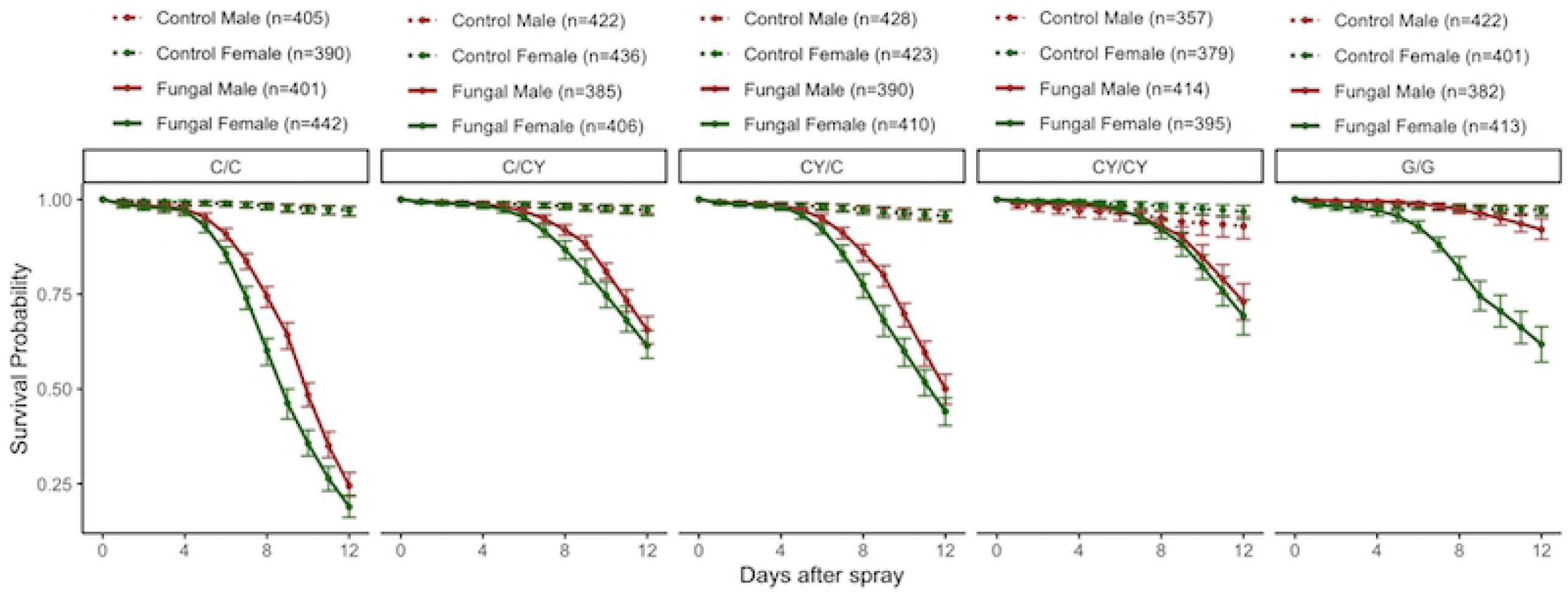
Yeast supplementation affects sexual dimorphism in survival of *D. melanogaster* inoculated with *B. bassiana* strain GHA. Data from Experiment 4. The figure shows model estimates for the raw data shown in Figure S4. Three days prior to control spray (dashed lines) or fungal spray (solid lines), flies were given Cornmeal (C), Cornmeal with yeast supplement (CY), or Glucose (G) diets (marked before the dash in the figure headings). After the spray, which was on day 15 from egg, flies were kept on one of the three types of diets (C, CY, or G, marked after the dash in the figure headings). Yeast supplementation improved survival of inoculated flies. When yeast supplementation was provided after the inoculation, sexual dimorphism was ablated. But there was sexual dimorphism among control flies when yeast supplementation was provided both before and after the spray. See Tables S6 for analysis of these results.

### The amount of supplemental yeast added to the diet impacts survival post inoculation with *B*. *bassiana* GHA (Experiment 5)

Control flies survived better than inoculated flies regardless of diet and did not present sexual dimorphism (Fig 6). The addition of yeast supplement after inoculation, when done at the same dose as in Experiment 4 (1 g yeast/ 5mL DI water) improved survival of both males and females relative to flies maintained on Cornmeal diets (Fig 6, Table S11) similar to the results of Experiment 4. At half the amount of yeast supplement, post inoculation survival was still improved relative to the Cornmeal-only diet, but survival was still significantly lower than with the higher dose of yeast supplementation (Table S11). When 1.5x the amount of yeast supplement was added relative to the amount in Experiment 4, the impact on post inoculation survival did not change (Table S11). Sexual dimorphism in survival post-inoculation was present in the 0-5 and 5-8 day intervals for the Cornmeal-only and lowest level of yeast supplementation (Table S12), but the sexual dimorphism was ablated in the 0-5 day interval at higher levels of yeast supplementation (Table S12).

**Fig 6:**
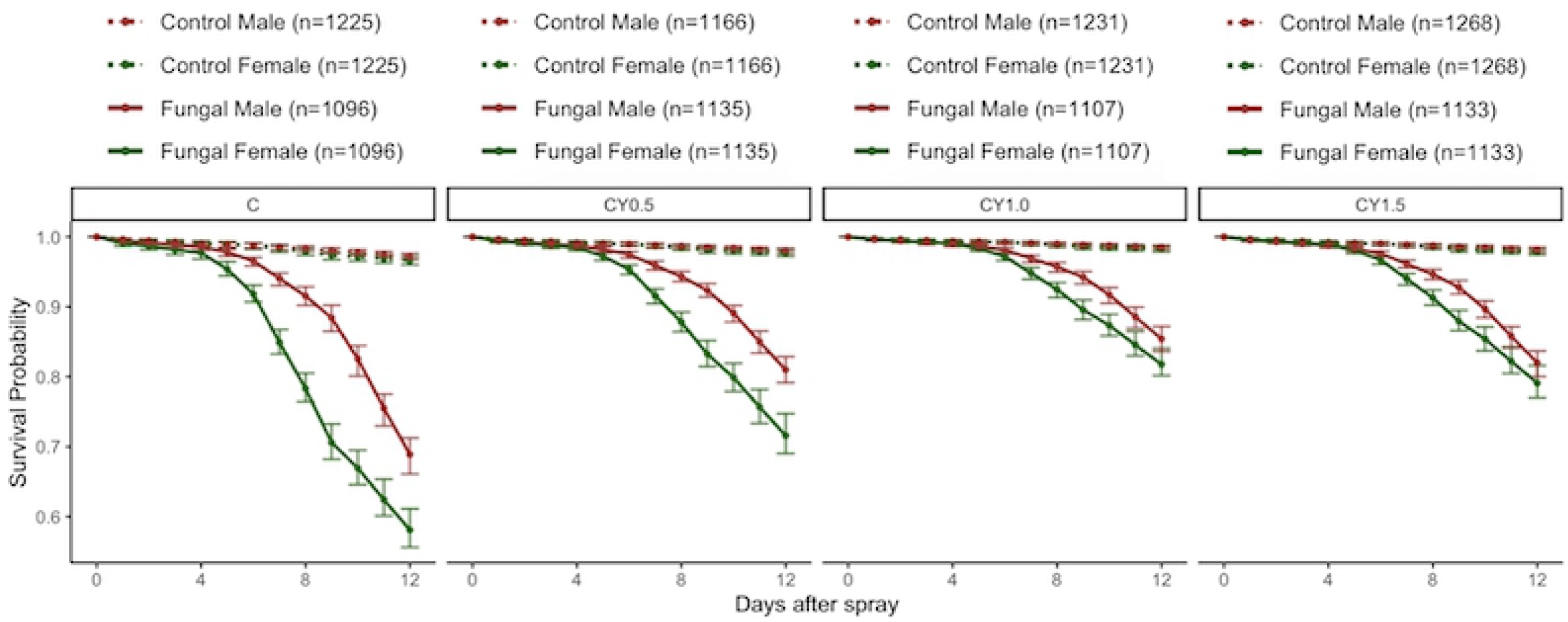
Level of yeast supplementation affects sexual dimorphism in survival when *D. melanogaster* are inoculated with *B. bassiana* GHA. The figure shows model estimates and 95% confidence intervals for the raw data shown in Figure S5. All flies were reared on cornmeal diets. After the sprays, flies received cornmeal diets supplemented with varying levels of yeast. See Tables S7 and S8 for statistical analysis of these results.

## Discussion

Shahrestani et.al. (11) found that *D. melanogaster* females from the outbred Population C3 were more susceptible than males to infection with *B. bassiana* strain GHA. Their study followed survival for ten days post inoculation and used cohabiting flies. To expand further on their result, we repeated their study but followed survival for 21 days post inoculation. While we replicated the results of Shahrestani et al. (11), we additionally found that with this fungal pathogen, sexual dimorphism in defense began to disappear around 12 days post inoculation, such that females and males had the same probability of survival from days 12-21 post inoculation. We also tested an additional *B. bassiana* strain, ARSEF 12460, that was shown to result in sexual dimorphism in defense in Shahrestani et al. (2018) with females from the Canton S population being more susceptible than males.

In our study with ARSEF 12460 the sexual dimorphism was reversed, such that males from Population C3 were more susceptible to fungal infection than females, and this dimorphism persisted through the 21 days of the experiment. One key difference between our study and Shahrestani et al. (11) is that they used the fly line Canton S to test susceptibility to ARSEF 12460, whereas we used the outbred Population C3. This suggests, that the host genotype influences the direction of sexual dimorphism in immune defense, and corroborates the result of Shahrestani et al. 2021 which showed that in ∼300 recombinant inbred fly lines the direction and magnitude of sexual dimorphism in immune defense was genotypespecific. Moreover, our result that in Population C3, the direction of sexual dimorphism was reversed when changing the fungal pathogen, suggests that sexual dimorphism in defense is also affected by the fungal strain used. Hence, there is no weaker sex when it comes to immune defense, and multiple factors, at least the pathogen and host genotypes, affect the direction of sexual dimorphism in defense.

Another variable that has been previously shown to affect sexual dimorphism in immune defense against bacterial pathogens is mating (12). Mating was shown to suppress defense in female *D. melanogaster*, such that mated females survived infection with *Providencia rettgeri* and *Providencia alcalifaciens* less successfully than virgin females (12). This mating-immunity trade-off has been thought of as a female trait (13) and it is generally assumed that males do not experience the same mating-immunity trade-off, due to sex differences in resource allocation to reproduction and defense. It is commonly expected that females will exert more energy during reproduction, thus producing an energy trade-off between reproduction and immune defense (12, 27). Therefore, one hypothesis for mating-immunity trade-offs is that female defense is suppressed when exposed to the male ejaculate accessory gland proteins (Acp) or sex peptide (SP) (14), due to the effects of juvenile hormone (24). However, reproduction in male *D. melanogaster* also accrues costs (29), and it has been suggested that these costs trade-off with immune defense (15). We therefore expected that a mating-immunity trade-off would be present for both males and females, and that these trade-offs could influence the direction of sexual dimorphism in defense. For both males and females, we tested survival after inoculation with *B. bassiana* ARSEF 12460 for virgin flies, flies that mated for 24 hours before inoculation, and flies that mated both before and after inoculation (males and females that cohabited together).

We found that both females and males experienced mating-immunity trade-offs. In both sexes, mating for 24-hours prior to inoculation made flies more susceptible to inoculation with ARSEF 12460 compared to virgins, and flies that cohabited with the other sex were more susceptible to inoculation than flies that only mated for one day. It is possible that the cause of the mating-immunity trade-off may be different for females and males. For females, we tested for differences in resource allocation by counting the number of offspring produced by females in different mating and infection conditions. Infected females produced fewer offspring than control females, suggesting that there is a reproductive cost to infection. Under uninfected conditions, females that cohabited with males produced more offspring than females that mated only for one day. However, under infected conditions, cohabiting and one-day mated females laid indistinguishable numbers of eggs, which may suggest that the increased susceptibility of cohabiting females was not due to increased egg laying. But it is also possible that, due to infection, the numbers of eggs laid were too low for differences to be picked up with our statistical analyses. We did not measure male reproductive output. However, other factors, outside of resource allocation to reproduction could potentially affect a trade-off with immune defense. In particular, mating itself is energetically costly, as is intrasex competition for mates. In our study, cohabiting flies were maintained in population cages with equal sex ratios. The stress and investment for successful mating could have affected male susceptibility to infection. Note that the fungus ARSEF 12460 is not itself transmitted by mating.

Interestingly, the direction of sexual dimorphism in defense against ARSEF 12460 varied by mating status. Among cohabiting flies, males were more susceptible to infection than females. However, when flies were mated for only 24-hours prior to inoculation, females were more susceptible to infection than males. Yet when virgin flies were compared, males were again more susceptible than females. This result supports the idea that the mating-immunity trade-off is at least in part affected by the frequency of mating, instead of being an inherent component of male and female biology.

Given the potential impacts of reproduction-related resource allocation on the mating-immunity trade-off and on sexual dimorphism in defense, our next avenue of research was to investigate the effects of dietary manipulations on these relationships. It is thought that diet affects mating preferences and survival in female and male fruit flies. For example, under uninfected conditions, mated females survive better on a more balanced diet compared to virgin females which survive better on more carbohydrate rich diets (25). The fecundity of mated females increases with increased ingestion of proteins (25). In a high energy and high carbohydrate diet there is a reduction in *D. melanogaster* eclosion time and adult survival (26). Egg to adult development time is faster in more protein rich diets than carbohydrate rich diets (27). In males, a high carbohydrate and low protein diet increases lifespan and reproductive performance (28), and in females, a high fat diet reduces lifespan (29) and mating (30). Female egg production is higher in protein rich diets compared to carbohydrate rich diets, and males survive better than females in both diets (31). Here we were specifically interested in determining the effects of diet on sexual dimorphism in survival after pathogen exposure.

There is some information about the effects of diet on *D. melanogaster* defense against bacterial pathogens. For example, a high sugar diet was seen to decrease immune defense against infection with *Providencia rettgeri*, resulting in decreased survival (32). To start to examine the effects of diet on survival of fungal infection, we used two diets that Population C3 had received in its evolutionary history, which differ drastically in their sugar and protein contents: the Cornmeal and Glucose diets (ingredients above). Historically at Cornell University, Population C3 was evolved on the Glucose diet where equal amounts of glucose and yeast were used for food preparation. However, when brought to the California State University in Fullerton, the Population C3 diet was switched to Cornmeal, which has a 6:1 ratio of cornmeal to yeast and a 3:1 ratio of molasses to yeast. Comparatively, the overall protein to carbohydrate ratio for the Glucose diet is 1:4, whereas Cornmeal is 1:9, as calculated by the Drosophila Dietary Composition Calculator (33). When comparing the effects of diet on immune defense, we learned that flies on their ancestral Glucose diet survived better than those on their more recent Cornmeal diet. Their survival may be due to them being more adapted to the Glucose diet, and/or it may be affected by the actual dietary contents.

Diet is known to affect *D. melanogaster* survival. *D. melanogaster* exhibited greater survival under conditions of low protein to carbohydrate ratios when under dietary restriction (34). With increased protein consumption, there was an increase in fecundity and a decrease in survival (35). In contrast, greater carbohydrate consumption led to lower fecundity and increased survival (35). However, in the presence of infection, flies displayed contrasting results. When infected with the bacterium Micrococcus luteus, flies survived better under diets of high protein to carbohydrate ratios (35). This may be indicative of the insect’s ability to selectively balance their nutrient intake and allocation based on their needs for infection resistance, survival and reproduction (36).

To test the effects of dietary contents, we explored the addition of supplemental yeast to the Cornmeal diet to determine if increased yeast improved the survival of the flies. The yeast supplementation provided additional protein to the surface of the Cornmeal diet in the form of a thin paste. The yeast supplementation consists of a 1:1 ratio of protein to carbohydrates (33) that offers an alternative and more balanced food source for the flies. Yeast supplementation improved the survival of both males and females under inoculated conditions and partially ablated sexual dimorphism in defense. This is in contrast to the Glucose diet, which while it also improved survival under inoculated conditions, did not get rid of sexual dimorphism in immune defense. Moreover, the timing of introduction of yeast mattered, with yeast provided after infection having greater benefits for survival of the infection than yeast provided before infection. Even low amounts of yeast supplementation improved survival of infection. Increasing the amount of yeast led to even greater benefits for survival but only up to intermediate yeast levels, beyond which further increases did not lead to additional benefits.

Survival of infection and sexual dimorphism in survival are complex and influenced many factors. We show that these factors include fly genotype, pathogen genotype, mating status, and diet. While we do not provide an extensive list of factors that influence postinfection survival, nor get into the mechanisms for how post-infection survival is affected, we believe that demonstrating the complexity in this trait, especially in sexual dimorphism of this trait, will help avoid the research pitfalls of looking for a weaker sex or treating mating-immunity trade-offs as a female-only phenomenon.

## Acknowledgements

This research was funded by a CSUPERB New Investigator Grant to PS. KER and PS conceived the research questions. KER, ALCB, MC, and PS devised experiments. KER, ALCB, MC, RC, and EB collected data. HY performed the analyses with guidance from SB, RR, and PS. KER, ALCB, MC, and PS prepared the manuscript. Several other students from the Shahrestani lab helped with data collection.

## Supporting Information

## Supplemental Figure Legends

**Fig S1. Sexual dimorphism in survival of D. melanogaster inoculated with B. bassiana strain ARSEF 12460 is affected by mating status**.

Data from Experiment 1. Female (green) and male (red) survival after control spray (dashed lines) and fungal spray (solid lines) is shown for cohabiting flies, virgin flies, and mated flies which mated for only 24 hours. Survival was followed for 21 days after the spray. Sample sizes per treatment are provided in the legend. The top graphs show model estimates for survival proportions, using four replicates of raw data with 95% Bootstrap confidence intervals. See Tables S1 and S2 for statistical analysis of this data. Bottom graphs show the raw data for the four replicates of each treatment and the means. For cohabiting and virgin flies, females had better survival than males after inoculation. For mated flies, this trend was reversed. In both females and males, virgin survival was higher than mated survival, which was itself higher than survival under cohabiting conditions.

## Supplemental Tables

**Table S1. Dietary treatments of Experiment 3**

For each dietary condition, half of the flies were inoculated and half were treated as controls. All flies were reared on a Cornmeal diet until age 12 from egg. Then some flies received Cornmeal and some received a Glucose diet. After flies were sprayed at age 15 days from egg, some flies received Cornmeal and some Glucose diet. Days here are given from eggs.

**Table S2. Dietary treatments of Experiment 4**

For each dietary condition, half of the flies were inoculated and half were treated as controls. All flies were reared on a Cornmeal diet until age 12 from egg. Then the specific dietary conditions were applied. Days here are given from egg.

**Table S3. Hazard ratios and *p*-Values when comparing mating statuses in both control and fungal inoculated *Drosophila melanogaster***

Data from Experiment 1.

Hazard ratios and *p*-Values are presented in the time intervals of 0-5, 5-11, and 11-21 days post inoculation.

**Table S4. Hazard ratios and *p*-Values when comparing males and females under different mating statuses in both control and fungal inoculated *Drosophila melanogaster***.

Data from Experiment 1. Hazard ratios and *p*-values are presented for days 0-21 post inoculation.

**Table S5. Table representing the interaction between days and treatment between the Control and Fungal inoculated reproductive output of the flies**.

Significantly smaller *p*-value of Treatment (2.505×10^−9^) indicates that the data strongly supports the difference on offspring counts between the fungal inoculated groups and the controls. The data also strongly supports that cohabiting and mated groups differ in their offspring counts (*p*-value: 1.564×10^−6^).

* p-value ≤ 0.05, *** p-value ≤ 0.001

**Table S6. Hazard ratios and *p*-values for males vs females in GHA inoculated flies** Data from Experiment 2. The hazard ratios indicate the risk of male flies dying post inoculation in comparison to the female flies. The data strongly supports that there is sexual dimorphism in post inoculation survival in the 0-8 and 8-12 age intervals. However the *p*value in the 12-21 days interval suggests there is no sexual dimorphism in survival in this interval.

**Table S7. Diet affects sexual dimorphism of *D. melanogaster* inoculated with *B. bassiana* strain GHA in a sex-specific manner**.

Data from Experiment 3. There was no sexual dimorphism in survival among the control treatments, regardless of diet. However, among fungal inoculated flies, males survived better than females in some age intervals on every diet. The timing of introduction of a Glucose diet has an impact on the Hazard ratio among inoculated flies.

Statistically significant differences are in Bold.

**Table S8. Effects of dietary condition and infection status on male and female survival in different age groups**

Data from Experiment 3.

**Table S9. Effect of diet on survival of males and females combined when inoculated with B. bassiana GHA**

Data from Experiment 4. Hazard ratios and *p*-values are presented from 0-4, 4-9 and 9-14 days post inoculation.

**Table S10. Yeast supplementation affects sexual dimorphism in surviving infection**. Data from Experiment 4. Whenever yeast supplementation was provided after inoculation, it ablated the sexual dimorphism in survival. However, when yeast supplementation was provided before and after the spray, there was sexual dimorphism among uninfected control flies. The hazard ratio, showing difference in hazard between males and females is largest for flies that received the glucose diet.

**Table S11. Experiment 5. Intermediate levels of yeast supplementation improved**

**survival**.

Pairwise comparisons of survival are shown, comparing the control (no yeast) condition with the lowest yeast level, the lowest and intermediate yeast levels, and the intermediate and high yeast levels. In both males and females, survival is improved by low and intermediate levels of yeast supplementation.

**Table S12. There was no sexual dimorphism in survival among control flies**.

Data from Experiment 5. When flies were inoculated with fungus, there was sexual dimorphism on all diets, but the age intervals and magnitudes of this dimorphism changed with the level of yeast supplementation. When there is no yeast supplement or a little amount of supplement, the dimorphism starts at earlier ages than with higher levels of yeast supplement.

